# A PCR-based assay for the sensitive and accurate determination of malaria parasite species that commonly infect humans, including discrimination of *Plasmodium ovalecurtisi* and *Plasmodium ovalewallikeri*

**DOI:** 10.64898/2026.10.21.760652

**Authors:** Faridah Ousseini, Mayumi Tachibana, Kongpope Chareonsuk, Kazuhide Yahata, Sheila Makiala-Mandanda, Yannick Munyeku-Bazitama, Richard Culleton

## Abstract

There are six *Plasmodium* species that commonly infect humans including *Plasmodium ovalecurtisi*, and *Plasmodium ovalewallikeri*. To ensure successful malaria control, it is important to track their prevalence and understand their epidemiological and clinical characteristics. Current PCR-based malaria diagnostic techniques targeting the highly sensitive mitochondrial genome do not distinguish between the two *Plasmodium ovale* species. Here we present a PCR-based species typing method targeting the mitochondrial COXIII involving a sensitive, nested PCR with species-specific primers and a simple enzyme digestion to distinguish *P. ovalecurtisi* and *P. ovalewallikeri*. The assay is highly specific to each species and highly sensitive, detecting 0.125 and 0.25 parasites per 1 µl of blood from *in vitro* cultured *Plasmodium knowlesi* and *Plasmodium falciparum* respectively. Furthermore, using 30 field samples, we validate its effectiveness in discriminating the two *P. ovale* spp., identifying 18 *P. ovalecurtisi*, 16 *P. ovalewallikeri*, 23 *P. falciparum* and 17 *Plasmodium malariae* infections. This novel assay allows the sensitive and accurate diagnosis of *P. falciparum*, *Plasmodium vivax*, *P. malariae*, *P. knowlesi*, *P. ovalecurtisi*, and *P. ovalewallikeri* and can be used to monitor human and zoonotic malaria infections for both diagnosis and epidemiological studies.

## 1. Introduction

Malaria may be caused by infection with one or more of numerous parasites of the genus *Plasmodium*. Ascertaining which species causes a particular infection is important for appropriate treatment and for epidemiological studies of parasite-species prevalence. In recent years, advances in molecular techniques have drastically improved malaria diagnosis, enhancing both the sensitivity of parasite detection and the specificity of species identification (Fitri et al., 2022). The common human-infecting *Plasmodium* species are *Plasmodium falciparum*, *Plasmodium vivax*, *Plasmodium malariae*, *Plasmodium ovalecurtisi and Plasmodium ovalewallikeri,* the latter two species having previously been considered one species, *Plasmodium ovale*, first described in 1922 (Stephens, 1922). In 1995, two morphologically identical variants of this species were described (Li et al., 1995) and in 2010, it was proposed that the two variants were in fact two distinct non-recombining species (Sutherland et al., 2010). The two species were initially given the trinomial names *P. ovale curtisi* and *P. ovale wallikeri* (Sutherland et al., 2010). As they are now thought to be species, not subspecies, they were renamed *P. ovalecurtisi* and *P. ovalewallikeri* in 2024 (Snounou et al., 2024), and we adopt this updated nomenclature here.

*Plasmodium ovale* spp. are generally less pathogenic than *P. falciparum* and *P. vivax,* causing milder symptoms and often remaining asymptomatic (Mueller et al., 2007). While remission is possible without treatment, *P. ovale* spp. infection can persist at low parasitaemia for extended periods of time. Additionally, both relapses (these species, like *P. vivax*, produce hypnozoites) and severe infections have been described (Strydom et al., 2014; Groger et al., 2019; Kotepui et al., 2020). There is some evidence that the two species may result in differing pathology (Rojo-Marcos et al., 2018; Mahittikorn et al., 2021b). They are often found in mixed infections with other *Plasmodium spp.* and current evidence suggests their prevalence was previously underestimated (Oguike et al., 2011; Yman et al., 2019; Akala et al., 2021; Abdulraheem et al., 2022).

In addition to the five species above, several zoonotic malaria species, including *Plasmodium knowlesi*, are now known to infect humans naturally (Mewara et al., 2023). First described in 1932 (Knowles & Gupta, 1932), *P. knowlesi* was long thought to be restricted to its natural non-human primate hosts. That it is a significant cause of malaria in humans was first shown in 2004, when many people were diagnosed with infections of this species in Sarawak, Malaysian Borneo (Singh et al., 2004). To date, evidence for human-to-human transmission of *P. knowlesi* remains limited, with most human infections likely occurring through bites from mosquitoes that previously fed on macaques (Fornace et al., 2023). However, the increasing number of cases threatens malaria control in affected regions, particularly Malaysia (Chin et al., 2020). In contrast to the erythrocytic replication cycle of 48 hours in *P. vivax*, *P. falciparum*, *P. ovale* spp. and 72 hours in *P. malariae*, *P. knowlesi* has a cycle of 24 hours and can rapidly result in very high parasitaemia, leading to major pathologies and even death if not diagnosed early (Cox-Singh et al., 2010), making species-specific diagnosis crucial for appropriate treatment.

Accurate and sensitive identification of the diverse *Plasmodium* species infecting humans is necessary to control malaria. Microscopy has been and continues to be the principal method of diagnosis in many regions. Unlike *P. falciparum*, *P. vivax* and *P. malariae* which may be easily distinguished by microscopy, *P. ovalewallikeri* and *P. ovalecurtisi* share identical morphology, making differentiation by microscopy impossible (Snounou et al., 2024). Additionally, *Plasmodium knowlesi* may be easily mistaken for *P. malariae* or *P. falciparum* by microscopy (Singh et al., 2004; Barber et al., 2013). While Rapid Diagnostic Tests (RDTs) offer a faster and more user-friendly approach for the detection of *Plasmodium* parasites in blood, they are often unable to specifically diagnose *P. malariae*, *P. ovale* spp. and *P. knowlesi*, and distinguish *P. ovalecurtisi* from *P. ovalewallikeri* (Maltha et al., 2013).

Molecular detection techniques targeting parasite DNA offer a highly sensitive approach to parasite detection and identification (Fitri et al., 2022). In particular, a nested PCR assay targeting the 18S rRNA gene developed in 1993 (Snounou et al., 1993) revolutionised malaria diagnostics, increasing sensitivity and enabling accurate differentiation of *P. falciparum*, *P. vivax*, *P. malariae*, and *P. ovale* spp. and with a later modification, *P. knowlesi* (Singh et al., 2004). In addition to the 18S rRNA gene, several other genes such as reticulocyte-binding protein 2 (rbp2), tryptophan rich antigen (tra), and lactate dehydrogenase (ldh) have been utilised for the detection of malaria parasites using both conventional and unconventional PCR methods (Oguike et al., 2011; Fitri et al., 2022). While some of these protocols report improved detection of both *P. ovale* spp. (Miller et al., 2015; Akerele et al., 2017), others have been specifically developed to distinguish between *P. ovalecurtisi* and *P. ovalewallikeri* with various levels of sensitivity (Oguike et al., 2011; Bauffe et al., 2012; Calderaro et al., 2012; Fuehrer & Noedl, 2014; Potlapalli et al., 2023; He et al., 2024).

The mitochondrial genome offers a particularly attractive target for PCR-based diagnosis of malaria due to its high copy number per parasite compared to most nuclear genes, and several assays that utilise it have been developed (Isozumi et al., 2015; Echeverry et al., 2016; Hayashida et al., 2017). Despite their higher sensitivity, very few address *P. knowlesi* or differentiate between *P. ovalecurtisi* and *P. ovalewallikeri*.

Here, we describe and validate a molecular diagnostic method targeting the mitochondrial Cytochrome C Oxidase subunit III (COXIII) gene that accurately discriminates all six *Plasmodium* spp. that commonly infect humans and differentiates *P. ovalecurtisi* from *P. ovalewallikeri*. This method involves a nested PCR that targets the COXIII gene, which identifies *P. falciparum*, *P. vivax*, *P. malariae*, *P. knowlesi*, and *P. ovale* spp., followed by a simple enzyme digestion to distinguish *P. ovalecurtisi* from *P. ovalewallikeri*.

## 2. Materials and Methods

### 2.1. Sample collection

Thirty *P. ovale* spp. positive dried blood samples collected as part of a population-based cross-sectional study in 2021 in Matadi, Democratic Republic of the Congo (DRC), were selected for this study (Munyeku-Bazitama et al., 2023). These samples were identified as *P. ovale* spp. positive using previously published primers (Isozumi et al., 2015). Genomic DNA was extracted using the QIAamp® DNA Mini Kit (QIAGEN, Hilden, Germany) from dried blood spots (∼20 µl of blood) and was eluted in 50 µl of elution buffer and stored at -20°C.

### 2.2. Ethics clearance

The current study was reviewed and approved by the ethics committee of Ehime University (No 2607003, Ver. 10.8, 2026). The secondary use of 30 *P. ovale* spp. positive samples collected in DRC during a SARS-CoV-2 population-based cross-sectional survey in 2021, was approved by the ethics committee of the Kinshasa School of Public Health (ESP/CE/37B/2022). Written informed consent was obtained from participants.

### 2.3. Parasite culture and dilution

O+ human erythrocytes (Japanese Red Cross Society) were used to culture *P. falciparum* strain 3D7 and *P. knowlesi* strain A1-H.1 following previously described protocols (Trager & Jensen, 1976; Moon et al., 2013).

Sorbitol synchronization of *P. falciparum* was performed to isolate ring stage parasites as previously described (Lambros & Vanderberg, 1979). Percentage parasitaemia was determined by microscopy of thin blood film smears, counting > 4000 red blood cells (RBCs). Synchronized ring stages of *P. falciparum* and mixed stages of *P. knowlesi* were each diluted in 50% hematocrit RBCs to give the following number of parasites per µl: 100, 10, 1, 10^-1^, 10^-2^, 10^-3^, and 0.

Twenty-five µl of the above dilution series was spotted onto filter paper (ADVANTEC^®^ Co., Ltd, Japan) and air dried overnight. DNA was extracted from both the freshly prepared blood solutions and from the dried blood spots following the QIAamp® DNA Mini Kit (QIAGEN) protocols for fresh blood and dried blood spots. The DNA was eluted in 100 µl Elution Buffer. An additional two-fold dilution was performed from the extracted DNA to give the following number of parasites per µl of the original blood: 1, 0.5, 0.25, 0.125 and 0.0625. Eluate corresponding to 1 µl of original blood was used in the primary PCR reaction for all the dilutions.

### 2.4. Plasmid construction and cloning

Cytochrome C Oxidase subunit III (COXIII) gene plasmids were designed for each species to assess the specificity and sensitivity of our primers. The pEX-A2J2 COXIII plasmids of *P. ovalecurtisi* (pEX-A2J2 Poc COXIII) and *P. ovalewallikeri* (pEX-A2J2 Pow COXIII) containing an ampicillin resistance gene were synthesized (Eurofins Genomics Japan). *Plasmodium falciparum*, *P. malariae*, *P. vivax*, and *P. knowlesi* COXIII genes were cloned into the pEX-A2J2 vector using In-Fusion^®^ HD cloning kit (Takara Bio Inc., Japan) per the manufacturer’s instructions. The In-Fusion primers used were 5’- TCACGGCATCTCGAGCTCGCCATTTGATAGCGGTTAACC-3’ (forward) and 5’- CCACTGACTGGATCCGATCCATACAGTCCCAGCGACAGCGG-3’ (reverse) with XhoI and BamHI restriction sites flanking the edges of the inserted COXIII gene. DH5*α* competent cells were used for the transformation, and the transformed cells were cultured on LB medium with ampicillin for selection, then in liquid PlusGrow medium with ampicillin overnight. Plasmid DNA was purified using QIAprep Spin Miniprep Kit (QIAGEN) and quantified using Qubit version 2.0 (Invitrogen, Thermo Fisher Scientific, Carlsbad, CA, USA).

### 2.5. COXIII Primer Design

Our assay is based on the previously described COXIII nested PCR of Isozumi et al. (2015). We redesigned the primary PCR to amplify a larger primary genus-specific fragment (∼1100 bp compared to ∼940 bp). This modification allowed the design of new *P. ovale* spp. primers that amplify a fragment with a HindIII restriction site in *P. ovalecurtisi* that is not present in *P. ovalewallikeri* (**Fig. 1A**). Additionally, we designed new specific secondary primers for *P. vivax* and *P. knowlesi*, as the previously published primers for *P. vivax* cross-react with *P. knowlesi*.

**Figure 1:**
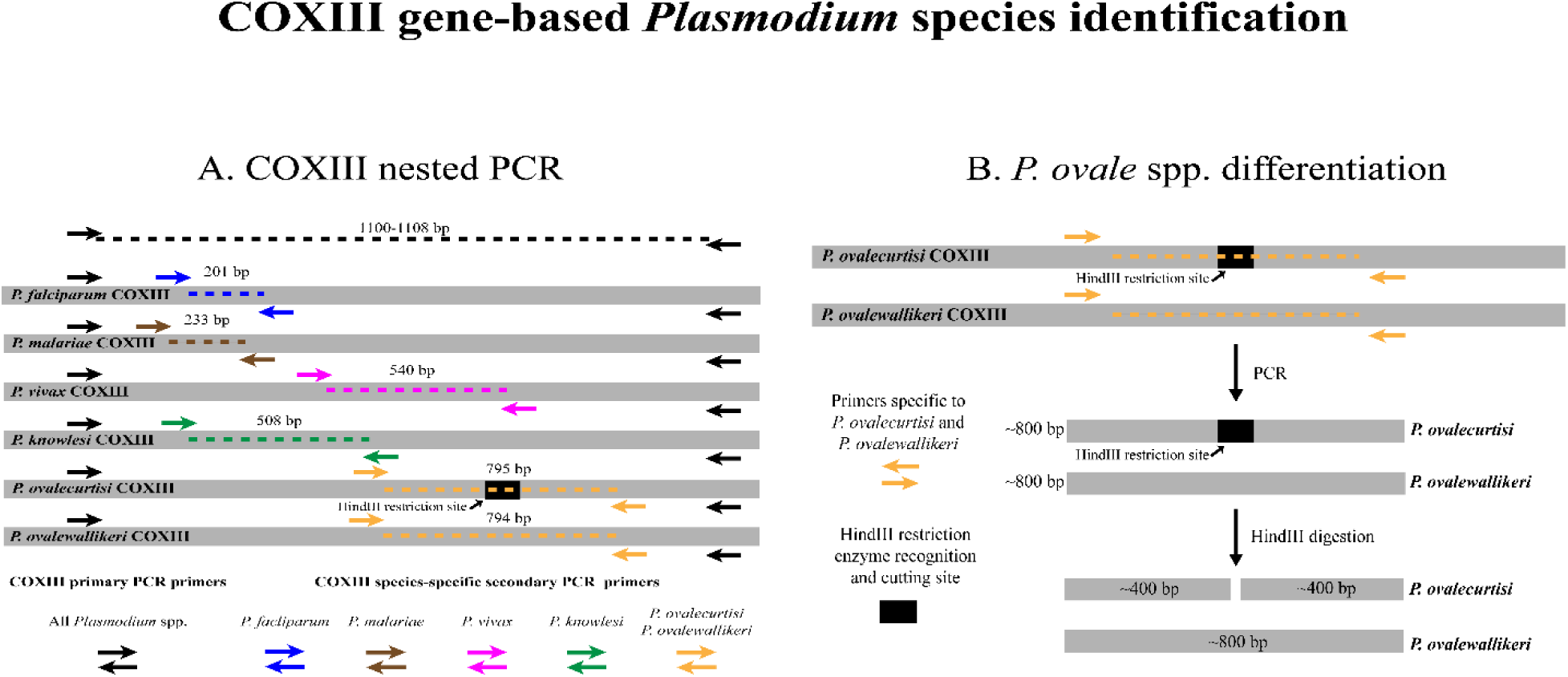
**A**: COXIII nested PCR design. **B**: *P. ovalecurtisi* and *P. ovalewallikeri* restriction enzyme differentiation assay based on a single HindIII restriction site.

The new primers (**Table 1**) were designed manually by aligning in MEGA version 11 (Tamura et al., 2021) the mitochondrial genome sequences of *P. ovalecurtisi* (GenBank accession number: HQ712052.1), *P. ovalewallikeri* (GenBank accession number: HQ712053.1), *P. falciparum* (GenBank accession number: JF923762.1), *P. malariae* (GenBank accession number: AB489194.1), *P. vivax* (GenBank accession number: KF668442.1), *P. knowlesi* (GenBank accession number: AY598141.1), and *P. cynomolgi* (GenBank accession number: KU245057.1).

**Table 1.**
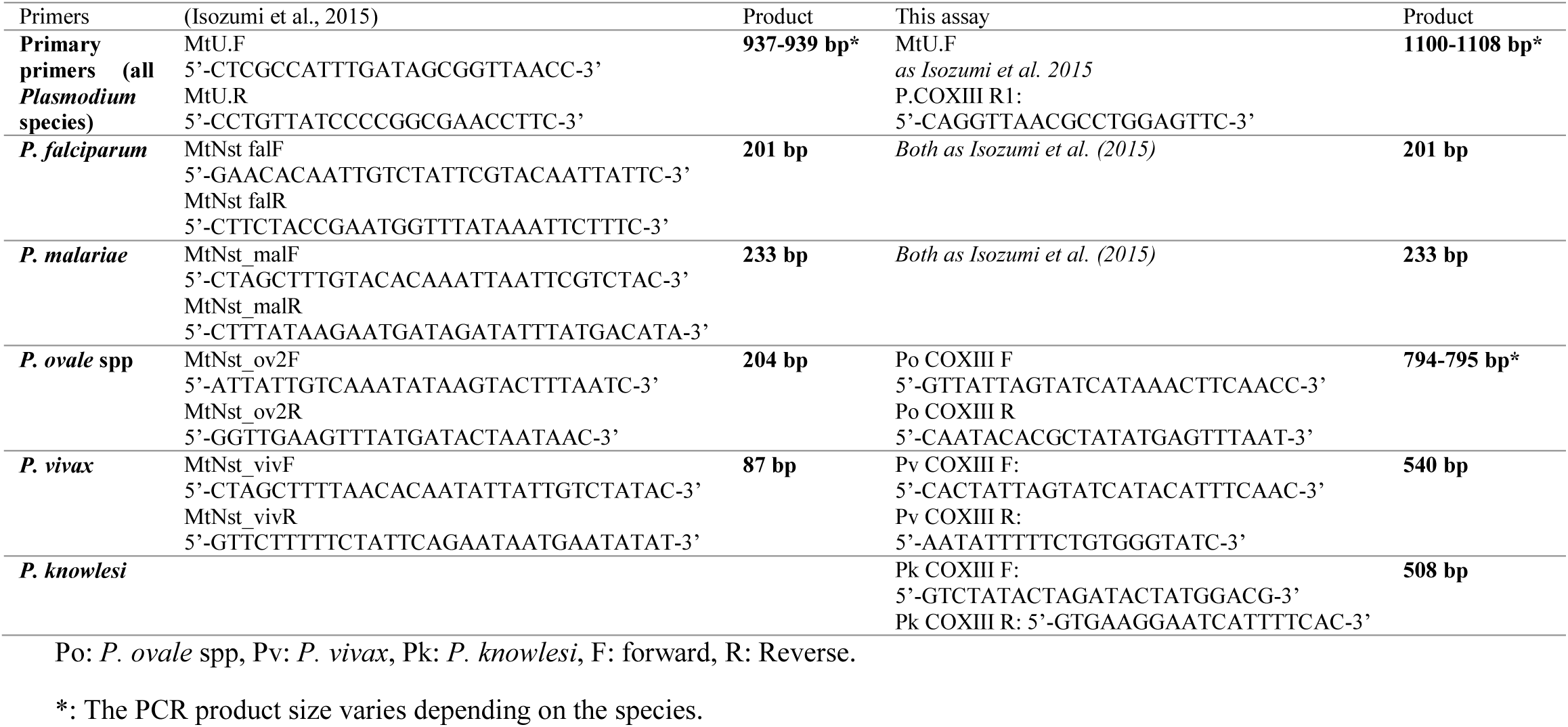
List of primary and secondary PCR primers.

Primer sequences and specificities were assessed with Primer BLAST (Ye et al., 2012) and their properties analysed with OligoAnalyzer (Integrated DNA Technologies, Inc., Coralville, IA, USA).

### 2.6. Nested Polymerase Chain Reaction

Newly designed primers were assessed for specificity using the *P. falciparum*, *P. malariae*, *P. vivax*, *P. ovalecurtisi*, *P. ovalewallikeri,* and *P. knowlesi* pEX-A2J2 COXIII plasmids. In the primary PCR, 12.5 µl of One*Taq*^®^ 2X Master Mix (New England Biolabs, Ipswich, MA, USA) and 0.4 µM of each primer were used in 25 µl reactions. Amplification was carried out under the following conditions: an initial denaturation at 94°C for 30 s, 40 cycles of denaturation at 94°C for 30 s, annealing at 59°C for 1 min, and extension at 68°C for 1 min 20 s, followed by a final extension at 68°C for 5 min.

The primary PCR product was diluted 100 times and used for the secondary PCR with each species-specific primer set to confirm the specificity of each of the new primer sets. The same primer set was used for *P. ovalecurtisi* and *P. ovalewallikeri*. One microliter of the diluted primary PCR product was used for the secondary PCR in a total 25 µl reaction containing 0.4 µM of each primer and 12.5 µl of One*Taq*^®^ 2X Master Mix (New England Biolabs). The amplification conditions were 94°C for 30 s for the initial denaturation, 35 cycles of 94°C for 30 s, 56°C for 1 min, 68°C for 45 s, and a final extension at 68°C for 5 min.

Additionally, we evaluated the primary PCR primers’ sensitivity using dilutions of *in vitro* cultured parasites following the same PCR protocol (diluting the primary PCR 50 times prior to secondary PCR) in comparison to the previously published COXIII primers (Isozumi et al., 2015). All PCR products were analysed on 1.5% agarose gels.

Finally, the new primary PCR primers along with the five sets of species-specific primers were used to screen the 30 field samples for *P. falciparum*, *P. malariae*, *P. vivax*, *P. ovalecurtisi*, *P. ovalewallikeri,* and *P. knowlesi*. Two microliters of the purified DNA (corresponding to approximately 0.8 µl of original blood) were used in a 10 µl primary PCR reaction and 2 µl of undiluted primary PCR product were used in a 50 µl secondary PCR following the above PCR conditions.

### 2.7. PCR purification

To distinguish between *P. ovalecurtisi* and *P. ovalewallikeri*, 45 µl of the secondary PCR products of *P. ovale* spp. positive samples were purified using QIAquick^®^ PCR Purification Kit (QIAGEN). The DNA was eluted in 50 µl of Elution Buffer.

### 2.8. Restriction Enzyme Assay

In PCR-based molecular detection assays for malaria parasites, *P. ovalecurtisi* and *P. ovalewallikeri* are often amplified with the same pair of primers due to their sequence similarity making them difficult to distinguish. Here, we developed a single restriction enzyme assay to distinguish the two following nested PCR amplification of the COXIII gene. The assay was designed to digest the secondary PCR product of *P. ovalecurtisi*, but not *P. ovalewallikeri*, using the restriction enzyme HindIII. The resulting two *P. ovalecurtisi* fragments (393 bp and 406 bp) appear as a single band of ∼400 bp and are half the length of *P. ovalewallikeri* (∼800 bp); thus, they may be distinguished following agarose gel electrophoresis (**Fig. 1B**).

In each reaction of 20 µl, ≤1 µg of the purified secondary PCR product was digested with 1 µl of HindIII (New England Biolabs) and 2 µl NEBuffer™ r2.1. The mixture was incubated at 37°C for 1 hour, and digested products run on 1.5% agarose gels.

### 2.9. Sequencing reaction

The purified secondary PCR products of *P. ovale* spp. positive samples were sequenced to confirm the results from the restriction assay. A total volume of 21 µl mixture containing 15-60 ng of purified secondary PCR product (794-5 bp), 12 µl of 0.8 µM primer (*P. ovale* spp. secondary PCR primers) and nuclease free water was prepared for each primer and sent for sequencing to Eurofins Genomics Japan. The sequence reads were aligned with the reference *P. ovalecurtisi* (GenBank accession number: HQ712052.1), *P. ovalewallikeri* (GenBank accession number: HQ712053.1), and analysed using ATCG software version 10.0.2 (GENETYX Corporation Japan 2021).

## 3. Results

### 3.1. The newly designed primers are species-specific

To test our new primers, we constructed COXIII gene plasmids for each of the six species (*P. falciparum*, *P. malariae*, *P. vivax*, *P. ovalecurtisi*, *P. ovalewallikeri,* and *P. knowlesi*). The primary PCR primers were selected to amplify all *Plasmodium* spp. and designed within the conserved regions of COXIII gene of *Plasmodium* species. As expected, we obtained an amplicon of around 1100 bp (**Fig. 2A**) from all six plasmids in the primary PCR reaction, confirming that the primers can amplify all six species.

**Figure 2:**
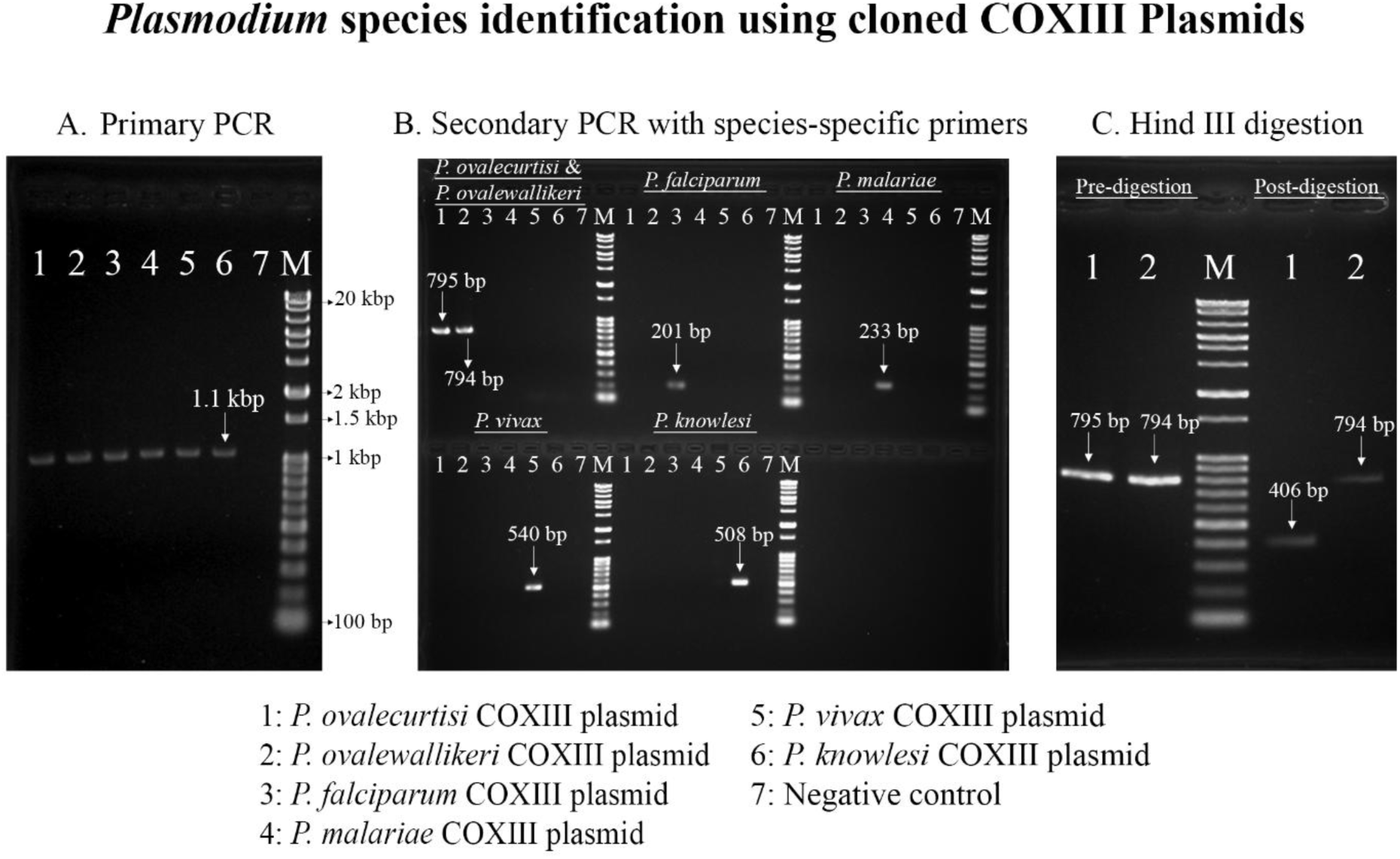
*Plasmodium* species identification using cloned *Plasmodium* COXIII plasmids. All samples were run on 1.5% agarose gels. **A**: Primary PCR with new generic *Plasmodium* spp. primers. **B**: Secondary PCR with species-specific primers using the primary PCR products as template. The same pair of primers was used for *P. ovalecurtisi* and *P. ovalewallikeri*. **C**: HindIII enzyme digestion of *P. ovalecurtisi* and *P. ovalewallikeri* secondary PCR products.

For the secondary PCR, we designed new sets of specific primers for *P. vivax*, *P. knowlesi*, and one set to amplify both *P. ovalecurtisi* and *P. ovalewallikeri* before differentiating them, aiming to reduce the difference in sensitivity between both *P. ovale* spp. as observed in other studies concerned with discrimination between the two (**Table 2**). For *P. falciparum* and *P. malariae*, we used the previously published COXIII species-specific primers (Isozumi et al., 2015). Initially, we included their *P. vivax* primers; however, as they cross-reacted with our *P. knowlesi* COXIII plasmid (data not shown), we designed new primers. We checked the specificity of each set of primers against the six species to ensure only the targeted species was amplified. The primary PCR product was used as a template for the secondary PCR, and five secondary PCR reactions were prepared separately for the five sets of primers. All the primers successfully amplified the corresponding species (**Fig. 2B**). The secondary PCR products obtained were 795 bp, 794 bp, 201 bp, 233 bp, 540 bp and 508 bp for *P. ovalecurtisi*, *P. ovalewallikeri*, *P. falciparum*, *P. malariae*, *P. vivax*, and *P. knowlesi*, respectively. The primer set for *P. ovalecurtisi* and *P. ovalewallikeri* amplified both as expected and did not amplify any other species. The same was observed with the other primer sets, confirming the specificity of each of them.

**Table 2.**
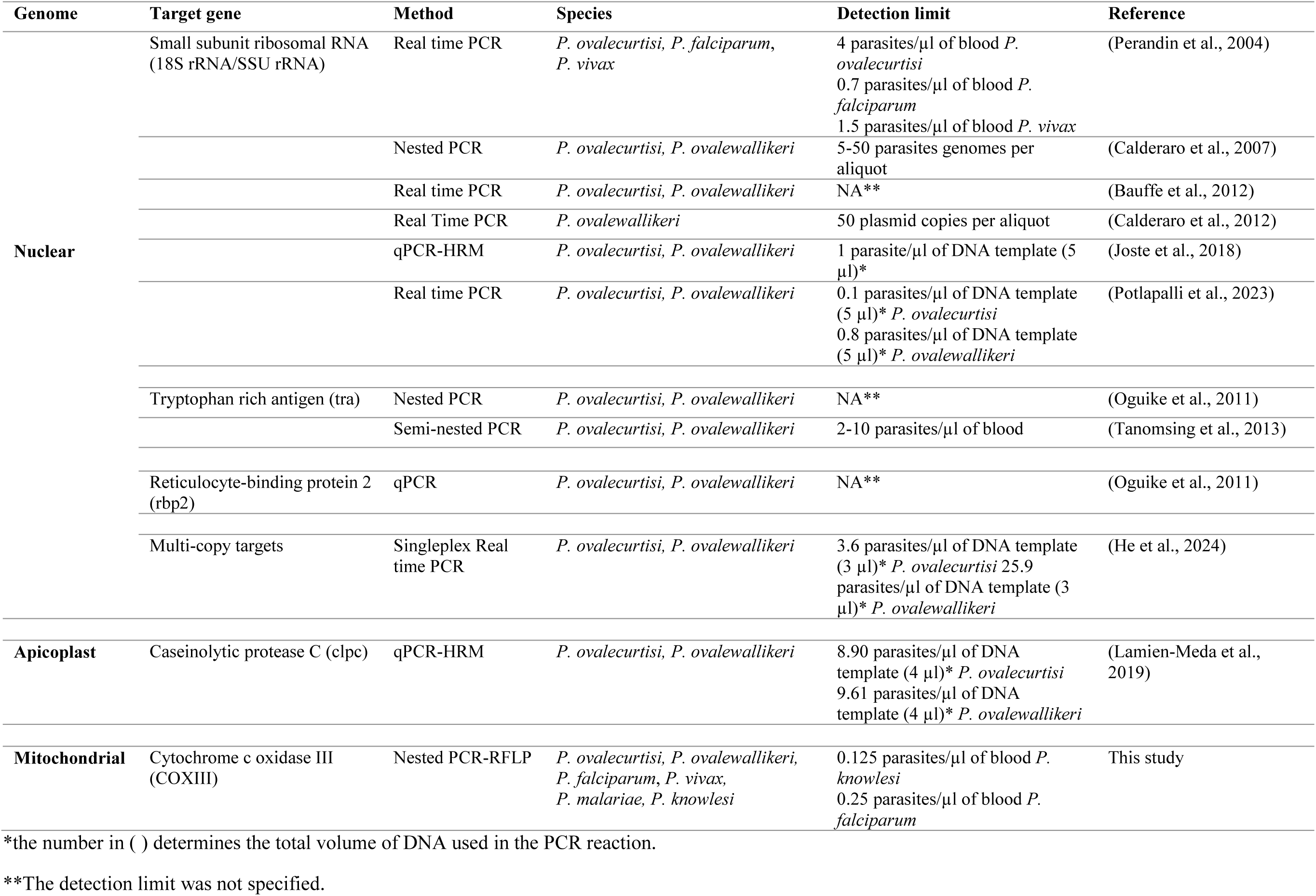
Molecular diagnosis assays for the discrimination *of Plasmodium ovalecurtisi* and *Plasmodium ovalewallikeri*.

Finally, to distinguish between *P. ovalecurtisi* and *P. ovalewallikeri*, the secondary PCR product was digested with HindIII restriction endonuclease. The primers specific for *P. ovalecurtisi* and *P. ovalewallikeri* were designed to produce an amplicon of about 800 bp with a HindIII restriction site in the middle of the *P. ovalecurtisi* COXIII amplicon which is absent in *P. ovalewallikeri*. The digestion resulted in two *P. ovalecurtisi* fragments seen as a single band of about 400 bp (**Fig. 2C**), while *P. ovalewallikeri* remained undigested (∼800 bp). Thus, we could differentiate between the two species without sequencing.

### 3.2. The assay can detect one parasite per PCR reaction

Next, we assessed the sensitivity of our primers in comparison to the previously published primers (Isozumi et al., 2015). We used *in vitro* cultured *P. falciparum* 3D7 and *P. knowlesi* A1-H.1 strains to determine the sensitivity of our assay. We performed ten-fold serial dilutions; from 100 to 10^-3^ parasites per µl (p/µl) of blood infected with synchronized ring stage *P. falciparum* parasites and asynchronous *P. knowlesi* parasites. The parasite numbers were determined from the % parasitaemia counted by microscopy per > 4000 RBC. Additionally, we compared the sensitivity of parasite DNA purified from fresh blood and dried blood spots, prepared as above. Each PCR reaction contained 4 µl of extracted DNA corresponding to 1 µl of original blood sample volume. From here on, we will describe detection sensitivity in terms of the number of parasites per µl (p/µl) of original blood sample. Each experiment was repeated at least twice. We did not assess the sensitivity of *P. malariae*, *P. vivax*, and *P. ovale* spp. primers due to a lack of an *in vitro* culture system and samples with known parasitaemias.

For *P. falciparum*, the primary PCR was able to detect 1 p/µl. No band was obtained in samples with ≤ 0.1 p/µl. Similarly, the secondary PCR was unable to detect samples ≤ 0.1 p/µl. Although the amplicon band appeared lighter in the PCR performed from DNA extracted from dried blood samples, no sensitivity difference was observed between the DNA samples derived from fresh and dried blood (**Fig. 3A**).

**Figure 3:**
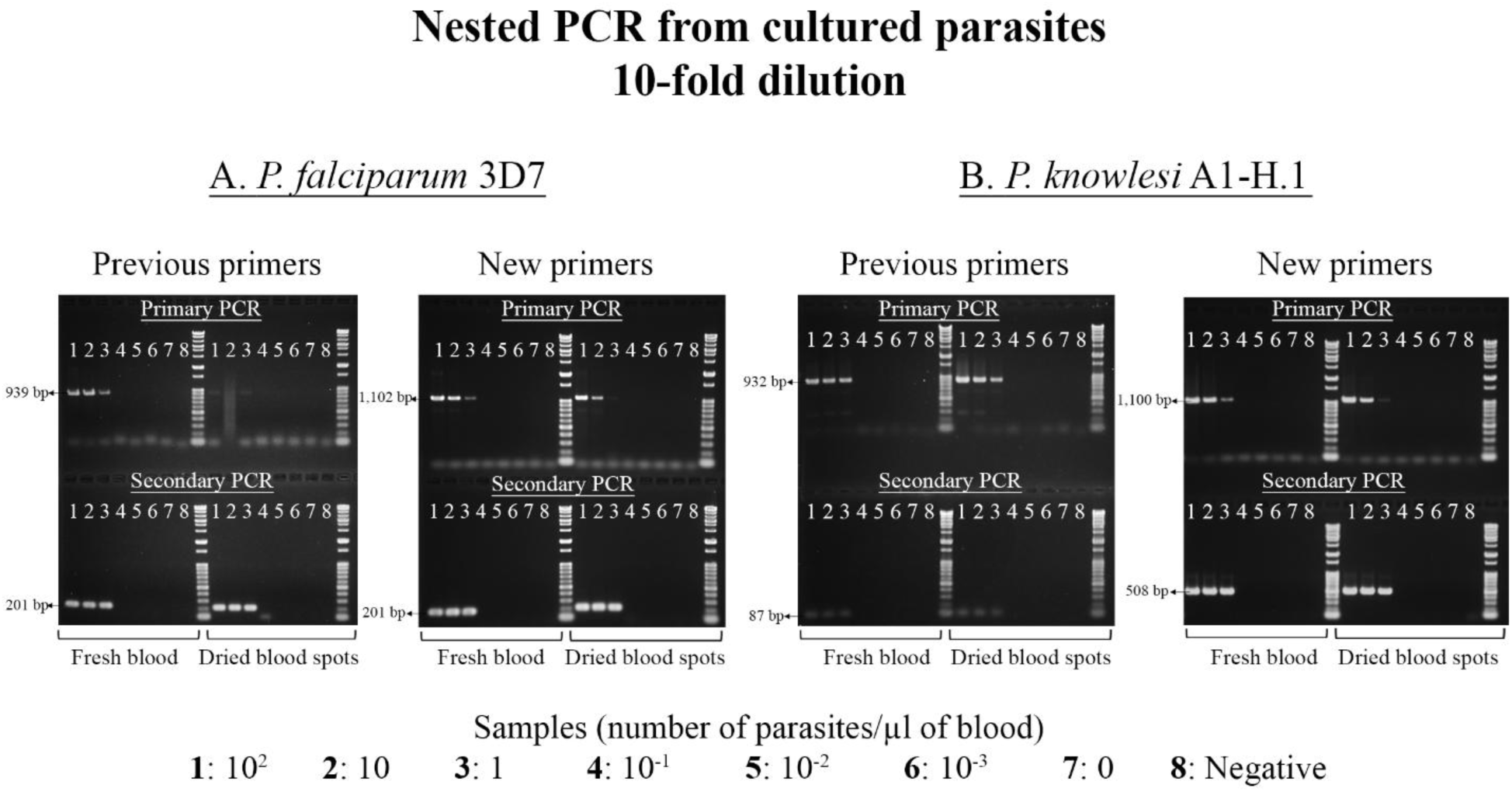
Primary and secondary PCRs of parasite dilutions (10-fold) using both the previously published primers (Isozumi et al., 2015) and the new primers; a comparison of their sensitivity in detecting DNA purified from fresh blood and dried blood spots. **A**: Nested PCR of *P. falciparum* parasite dilutions. **B**: Nested PCR of *P. knowlesi* parasite dilutions. *P. vivax* primers were used as the secondary PCR primers for *P. knowlesi* in the nested PCR with the previously published primers. The agarose gel concentration was 1.5% for all gels.

We obtained similar results with the previously published primary PCR primers (Isozumi et al., 2015) for the secondary PCR reaction. In the primary PCR reaction on the other hand, the bands were fainter in the dried blood samples compared to the fresh blood samples. Furthermore, a smear was produced in the 10 p/µl dried blood sample. No such smear is seen in the secondary PCR. The same was observed in repeated experiments (data not included). This could be due to the DNA quality or primer annealing conditions. However, when the same sample was amplified with the newly designed primers, the expected band was observed. It is important to note that we used the same *P. falciparum*-specific secondary primers published by Isozumi et al. (2015) in both experiments. The only difference is the primary PCR reverse primer, which was newly designed.

For the *P. knowlesi* assay, both the primary and secondary primers differed from those previously published. The Isozumi et al. (2015) protocol does not include *P. knowlesi* primers. However, as their *P. vivax* primers also amplify *P. knowlesi*, we used them here as secondary PCR primers to assess the new primers. Similar to what was observed for *P. falciparum*, a positive band was obtained in *P. knowlesi* samples with ≥1 p/µl but not in samples with ≤ 0.1 p/µl in the primary and secondary PCR (**Fig. 3B**). This is seen in the primary and secondary PCR from the new primers as well as the primers from Isozumi et al. (2015), although here, the bands from the primary PCR with the previous primers were brighter in the dried blood samples compared to the new primers, especially in the 1 p/µl samples. We also noticed a lighter unspecific band around 300 bp in the primary PCR with the previous primers. The primary PCR primers of Isozumi et al. (2015) were not tested on *P. knowlesi* in their study due to the lack of positive control DNA. Furthermore, similar faint smearing was observed in some other samples from both previous and new primers, especially in the primary PCR, which could be due to the shearing of DNA template or excessive DNA amount.

Additionally, even though we used synchronized ring stages of *P. falciparum* and asynchronous mixed stages of *P. knowlesi*, we did not see a difference in sensitivity between the two species (we assume that an asynchronous infection would contain relatively higher copy numbers of the mitochondrial genome per parasite than synchronous ring stage infection).

To determine the lowest detection limit, we prepared a two-fold dilution below 1 p/µl from the purified DNA and performed nested PCR. Contrary to the ten-fold dilution, we observed higher sensitivity from the fresh blood samples compared to the dried blood samples in both species (**Fig. 4**).

**Figure 4:**
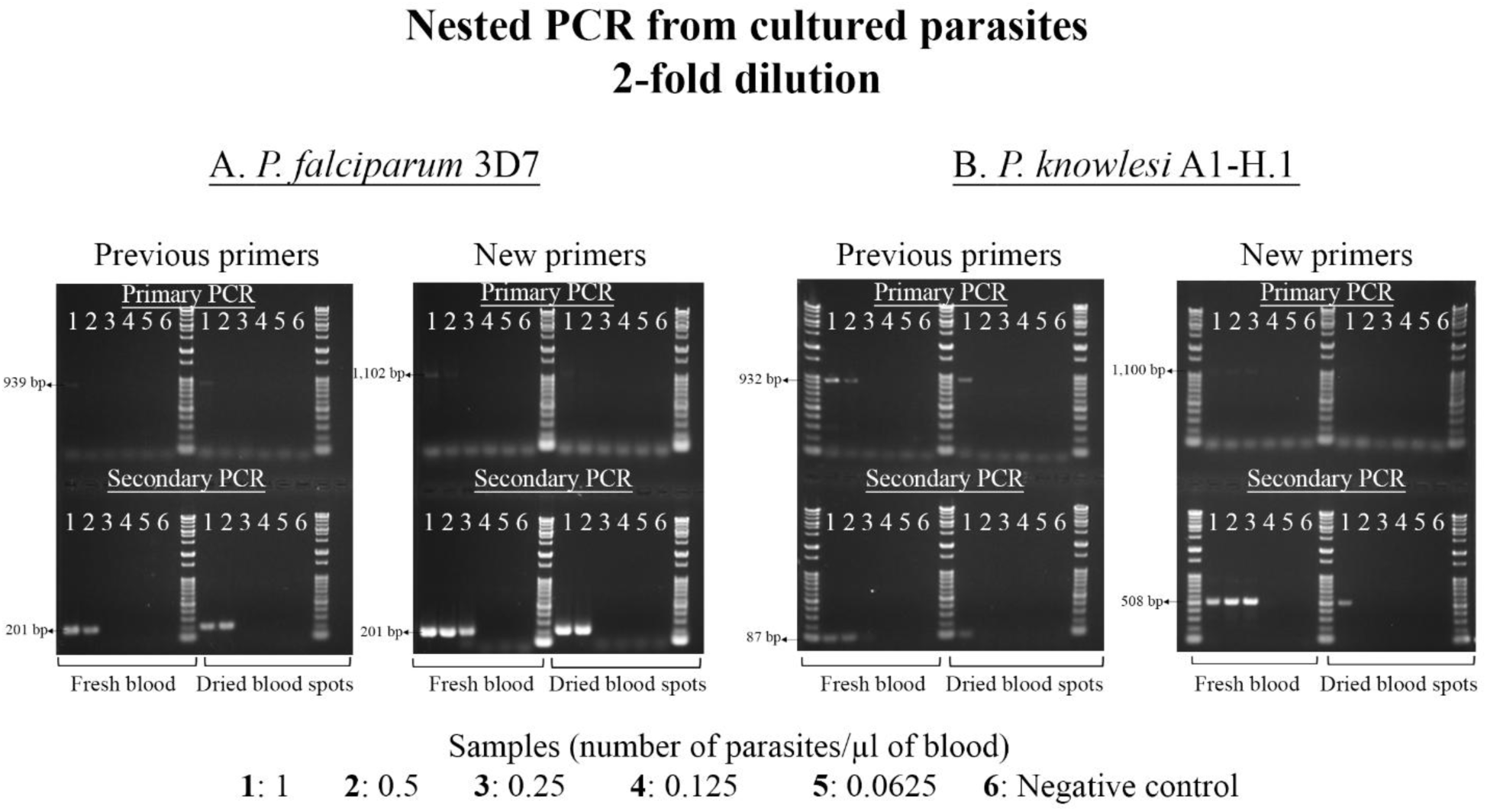
Primary and secondary PCRs of parasite dilutions (2-fold) using both the previously published primers (Isozumi et al., 2015) and the new primers; a comparison of their sensitivity in detecting DNA purified from fresh blood and dried blood spots. **A**: Nested PCR of *P. falciparum* parasites dilutions. **B**: Nested PCR of *P. knowlesi* parasites dilutions. *P. vivax* primers were used as the secondary PCR primers for *P. knowlesi* in the nested PCR with the previously published primers. The agarose gel concentration was 1.5% for all gels.

For *P. falciparum* (**Fig. 4A**), using the previous primers, we could detect 1 p/µl in the primary PCR and up to 0.5 p/µl in the secondary PCR from both fresh and dried blood samples. With the new primers, we could detect up to 0.5 p/µl and 0.25 p/µl from the fresh blood samples in the primary and secondary PCR, respectively. From the dried blood spot samples, the lowest detection was 1 p/µl in the primary PCR and 0.5 p/µl in the secondary PCR.

For *P. knowlesi* (**Fig. 4B**), the lowest detection from fresh blood samples was 0.5 p/µl and 0.25 p/µl in the primary and secondary PCR, respectively, using the previous primers. The limit was 1 p/µl from the dried blood samples in both primary and secondary PCR. Similarly, with the new primers, we could only detect 1 p/µl in the dried blood samples in both primary and secondary PCR. However, the lowest detection in the fresh blood samples was 0.25 p/µl in the primary PCR and 0.125 p/µl in the secondary PCR.

### 3.3. Identification of Plasmodium species in field samples from the DRC, including discrimination of P. ovalecurtisi and P. ovalewallikeri

To test our assay on field samples, we selected 30 *P. ovale* spp. positive samples from DRC (Munyeku-Bazitama et al., 2023), diagnosed as such using the primers described by Isozumi et al. (2015). We aimed to distinguish between *P. ovalecurtisi* and *P. ovalewallikeri* in addition to the other species using all five primer sets. The COXIII plasmids used to confirm primer specificity were used as positive controls.

In the primary PCR, 28 positive bands were observed using the new *Plasmodium* spp. generic primary PCR primers (**Fig. 5A**). In the secondary PCR reactions, using species-specific primers, 23 samples were positive for *P. falciparum*, 17 for *P. malariae,* and all 30 samples were positive for *P. ovale* spp. (**Fig. 5B**, **5C & 5D**), confirming the previous results. None of the samples were positive for *P. vivax* or *P. knowlesi* (**Fig. 5E & 5F**). These results indicate that the new primers are species-specific using natural infections sampled on dried blood spots.

**Figure 5:**
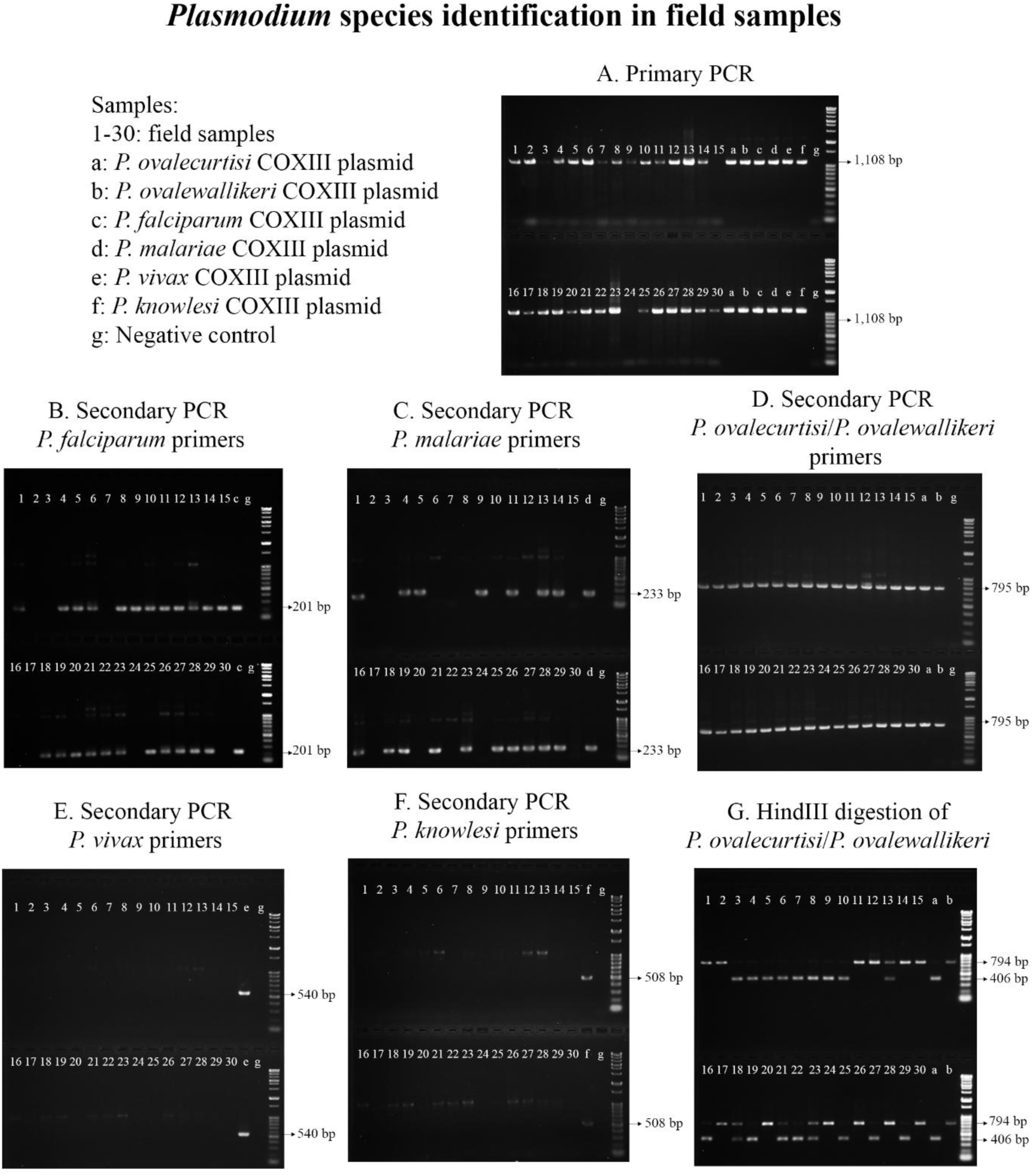
*Plasmodium spp.* identification using the new primers and restriction assay. **A**: Primary PCR of all 30 samples using COXIII plasmids as positive controls. **B-F**: Secondary PCR using species-specific primers and primary PCR products as template. **G**: HindIII digestion of *P. ovalecurtisi/P. ovalewallikeri* secondary PCR products discriminating between the two species. All samples were run on 1.5% agarose gels.

Next, to differentiate *P. ovalecurtisi* from *P. ovalewallikeri,* we purified the secondary PCR products of the *P. ovale* spp. samples and digested them with HindIII. **Fig. 5G** depicts a clear distinction between the two species based on their band size on an agarose gel. From the 30 samples, 18 contained *P. ovalecurtisi* (∼400 bp) and 16 contained *P. ovalewallikeri* (∼800 bp), with four containing both, indicated by double bands. While samples with a *P. ovalewallikeri* ratio lower or similar to *P. ovalecurtisi* were easier to identify (**Fig. 5G**, sample 13, 18 and 23), samples with a low *P. ovalecurtisi* to *P. ovalewallikeri* ratio could be confused with an incomplete digestion of *P. ovalewallikeri*. However, although the amount of *P. ovalecurtisi* was lower in sample 21 than *P. ovalewallikeri*, it was still distinguishable from single *P. ovalewallikeri* infections.

To validate the restriction assay, we sequenced the purified *P. ovale* spp. COXIII secondary PCR products and confirmed the species using the reference *P. ovalecurtisi* (GenBank accession number: HQ712052.1) and *P. ovalewallikeri* (GenBank accession number: HQ712053.1) sequences. The sequences matched the species identified using the restriction assay (**Table 3**). Additionally, one SNP m.248C>T (T83I) was observed between *P. ovalecurtisi* samples and the reference sequence. This SNP results in a change from threonine (T) to Isoleucine (I) at the 83^rd^ amino acid position of the enzyme. In *P. ovalewallikeri* samples, three SNPs m.16A>T (K6X), m.423C>T (L141L), and m.665G>A (G222E) were observed between the field samples and the reference sequence. The mutation m.16A>T is non-synonymous, and results in a change from a Lysine (K) in the reference genome to a potential stop codon (TAA) in the field samples. All the mutated positions had a single peak in the electropherogram. As the *P. ovalewallikeri* samples were processed in the order given in **Table 3** along with *P. ovalecurtisi* samples, we ruled out the possibility of mixed templates. Furthermore, we repeated the experiment from the original DNA samples to verify any sequencing artefact and confirmed the presence of the mutations.

**Table 3.** Polymorphisms in the partial COXIII amplicon of *P. ovalecurtisi* and *P. ovalewallikeri* from the DRC.

| Sample | HindIII results | Sequence | SNPs | Amino acid | Species present |
| --- | --- | --- | --- | --- | --- |
| 1 | POW | POW | 16A>T, 423C>T, 665G>A | K6X, L141L, G222E | POW+PM+PF+ |
| 2 | POW | POW | 16A>T, 423C>T, 665G>A | K6X, L141L, G222E | POW+ |
| 3 | POC | POC | 248C>T | T83I | POC+ |
| 4 | POC | POC | 248C>T | T83I | POC+PM+PF+ |
| 5 | POC | POC | 248C>T | T83I | POC+PM+PF+ |
| 6 | POC | POC | 248C>T | T83I | POC+PF+ |
| 7 | POC | POC | 248C>T | T83I | POC+ |
| 8 | POC | POC | 248C>T | T83I | POC+PF+ |
| 9 | POC | POC | 248C>T | T83I | POC+PM+PF+ |
| 10 | POC | POC | 248C>T | T83I | POC+PF+ |
| 11 | POW | POW | 16A>T, 423C>T, 665G>A | K6X, L141L, G222E | POW+PM+PF+ |
| 12 | POW | POW | 16A>T, 423C>T, 665G>A | K6X, L141L, G222E | POW+PF+ |
| 13 | POC+POW | POC+POW | * |  | POC+POW+PM+PF+ |
| 14 | POW | POW | 16A>T, 423C>T, 665G>A | K6X, L141L, G222E | POW+PM+PF+ |
| 15 | POW | POW | 16A>T, 423C>T, 665G>A | K6X, L141L, G222E | POW+PF+ |
| 16 | POC | POC | 248C>T | T83I | POC+PM+ |
| 17 | POW | POW | 16A>T, 423C>T, 665G>A | K6X, L141L, G222E | POW+ |
| 18 | POC+POW | POC+POW | * |  | POC+POW+PM+PF+ |
| 19 | POC | POC | 248C>T | T83I | POC+PM+PF+ |
| 20 | POW | POW | 16A>T, 423C>T, 665G>A | K6X, L141L, G222E | POW+PF+ |
| 21 | POC+POW | POC+POW | * |  | POC+POW+PM+PF+ |
| 22 | POC | POC | 248C>T | T83I | POC+PF+ |
| 23 | POC+POW | POC+POW | * |  | POC+POW+PM+PF+ |
| 24 | POW | POW | 16A>T, 423C>T, 665G>A | K6X, L141L, G222E | POW+ |
| 25 | POC | POC | 248C>T | T83I | POC+PM+PF+ |
| 26 | POW | POW | 16A>T, 423C>T, 665G>A | K6X, L141L, G222E | POW+PM+PF+ |
| 27 | POC | POC | 248C>T | T83I | POC+PM+PF+ |
| 28 | POW | POW | 16A>T, 423C>T, 665G>A | K6X, L141L, G222E | POW+PM+PF+ |
| 29 | POC | POC | 248C>T | T83I | POC+PM+PF+ |
| 30 | POW | POW | 16A>T, 423C>T, 665G>A | K6X, L141L, G222E | POW+ |

POC: *P. ovalecurtisi*, POW: *P. ovalewallikeri*, PM: *P. malariae*, PF: *P. falciparum*
\*The SNPs observed in POC and POW samples were also present in mixed POC+POW infections.

**Table 3** summarizes the species identified from the 30 samples used in this study. We obtained six single infections (two *P. ovalecurtisi* and four *P. ovalewallikeri*) and eight double infections comprising four *P. ovalecurtisi* + *P. falciparum*, one *P. ovalecurtisi* + *P. malariae*, and three *P. ovalewallikeri* and *P. falciparum*. The triple infections totalled 12, including seven *P. ovalecurtisi*+ *P. falciparum* + *P. malariae* and five *P. ovalewallikeri* + *P. falciparum* + *P. malariae*. Finally, the number of quadruple infections *P. ovalecurtisi* + *P. ovalewallikeri* + *P. falciparum* + *P. malariae* was four.

## 4. Discussion

The mitochondrial genome of the malaria parasite is multicopy, with previous estimates of its copy number ranging from 20-150 copies per parasite, depending on the species (Vaidya & Arasu, 1987; Joseph et al., 1989; Preiser et al., 1996). This is ∼ 4-20 times higher than the copy number of the 18S rRNA gene, which has been the standard target for malaria detection by PCR (Snounou et al., 1993). Several PCR diagnostic methods have been developed based on the mitochondrial genes Cytochrome b (Cyt b), Cytochrome C Oxidase subunit I (COXI), and COXIII (Costa et al., 2023; Deora et al., 2025), although they do not distinguish between the two *P. ovale* species. Here, we present a novel assay to diagnose the six most common *Plasmodium* species infecting humans, including *P. ovalecurtisi* and *P. ovalewallikeri* based on the COXIII gene.

This mitochondrial genome-based method accurately and sensitively distinguishes *P. ovalecurtisi* and *P. ovalewallikeri* from each other and from the four other common *Plasmodium* species, without the need for nucleotide sequencing of the PCR products. It is able to detect as low as 0.125 p/µl of fresh cultured blood (2.5×10^-6^ % parasitaemia) for *P. knowlesi* and 0.25 p/µl (5×10^-6^ % parasitaemia) for *P. falciparum*, when the initial PCR reaction contains DNA extracted from 1 µl of blood, a sensitivity higher than most assays reported to date (Deora et al., 2025).

Evaluating the difference in sensitivities between malaria parasite detection assays that use either fresh blood or dried blood on filter papers is difficult due to the differing volumes used to extract DNA from the two sources, with less total blood volume typically used for dried blood spots (Mahittikorn et al., 2021a). In this study, while we observed no difference in sensitivity between fresh blood and dried blood spot samples when using the same blood volume in higher parasitaemia samples (≥1 p/µl), we observed a higher sensitivity with fresh blood samples compared to dried blood samples in dilutions below 1 p/µl.

The COXIII gene is a structurally important component of the mitochondrial electron transport chain along with the two other mitochondrial genes, COXI and Cyt b (Hikosaka et al., 2011; Ramesh et al., 2023). Very few polymorphisms have been reported in the *Plasmodium* COXIII gene, in contrast to the *cytb* mutations reported to be involved in resistance to the antimalarial drug atovaquone (Syafruddin et al., 1999). We report polymorphisms in *P. ovalecurtisi* (T83I) and *P. ovalewallikeri* (K6X, L141L and G222E). The presence of the stop codon TAA at the sixth amino acid of the predicted COXIII enzyme in *P. ovalewallikeri* samples is surprising. Considering the necessity of the mitochondrial Electron Transport Chain to the viability of the parasite (Painter et al., 2007; Ke et al., 2011; Gujarati et al., 2026), it is very unlikely that COXIII gene results in a non-functional protein. We did not find this mutation in *P. ovalecurtisi* samples, nor was it present in *P. ovale* spp. COXIII sequences available online (GenBank HQ712052.1, HQ712053.1, AB354571.1, PP317149.1; PlasmoDB POWCR01_000243600, PocGH01_MIT000090). TAA is the most frequent stop codon in the *Plasmodium* mitochondrial genome and is specifically recognized by the release factor PfRF1Mit in *P. falciparum* to halt translation (Vaishya et al., 2016). While no other codon substitutions have been reported for TAA, internal TAA stop codon read-through has been reported in *P. falciparum* in the multigene family Pf60 expressed during the blood stages of the parasites (Bischoff et al., 2000). Another possibility is the presence of an alternative start codon downstream of this mutation.

Although the method presented in this study is PCR-based and may not provide instant results as required for diagnosis at the point of care in health care settings, it could be useful in epidemiological studies as it provides sensitive identification of all six human *Plasmodium* species, and removes the requirement for further sequencing to identify *P. ovale* spp.

We acknowledge that mutations in the highly specific HindIII restriction site could limit the distinction between *P. ovalecurtisi* and *P. ovalewallikeri* in the unlikely situation of a mutation at this nucleotide position. Furthermore, while single *P. ovale* spp. infections are easily identified in this assay, mixed *P. ovalewallikeri* and *ovalecurtisi* infections may be difficult to diagnose when *P. ovalecurtisi* is at very low parasitaemia compared to *P. ovalewallikeri*, as this could result in a result that is difficult to distinguish from incomplete enzyme digestion of single *P. ovalewallikeri* infection. However, our only sample having a lower ratio of *P. ovalecurtisi* compared to *P. ovalewallikeri* was clearly distinguishable from single *P. ovalewallikeri* infections. We were unable to assess the detection limit of this assay for *P. ovale* spp., *P. malariae*, and *P. vivax* due to a lack of samples with known parasitaemias.

In conclusion, the current study presents a new assay based on the mitochondrial COXIII gene that distinguishes all six common *Plasmodium* species that infect humans, including *P. ovalecurtisi* and *P. ovalewallikeri* with high sensitivity and specificity.

## Author contributions

**Faridah Ousseini:** Conceptualization, Investigation, Methodology, Writing-Original Draft. **Mayumi Tachibana:** Supervision, Resources, Writing-Review & Editing. **Kongpope Chareonsuk:** Resources, Investigation. **Kazuhide Yahata:** Supervision, Resources, Writing-Review & Editing. **Sheila Makiala-Mandanda**: Investigation, Resources, Writing-Review & editing. **Yannick Munyeku-Bazitama**: Investigation, Resources, Writing-Review & editing. **Richard Culleton:** Conceptualization, Methodology, Supervision, Writing-Review & Editing, Validation.

## Acknowledgements

The authors are grateful to the Japan International Cooperation Agency Ogata Sadako Research Institute for Peace and Development for the financial support provided under the COVID-19 research project, thanks to which clinical samples used in this study were collected. They also thank all study participants and the Division Provinciale de Santé du Kongo Central. The first author gratefully acknowledges the financial support provided by the Japanese Government (MEXT) Scholarship. Authors are also thankful to the Japanese Red Cross Society for providing human erythrocytes used in parasites *in vitro* culture. Special thanks to Aki Konishi and Sono Sadaoka for their technical support.

## References

Abdulraheem, M. A., Ernest, M., Ugwuanyi, I., Abkallo, H. M., Nishikawa, S., Adeleke, M., Orimadegun, A. E., & Culleton, R. (2022). High prevalence of *Plasmodium malariae* and *Plasmodium ovale* in co-infections with *Plasmodium falciparum* in asymptomatic malaria parasite carriers in southwestern Nigeria. Int. J. Parasitol., 52(1), 23–33. 10.1016/j.ijpara.2021.06.003

Akala, H. M., Watson, O. J., Mitei, K. K., Juma, D. W., Verity, R., Ingasia, L. A., Opot, B. H., Okoth, R. O., Chemwor, G. C., Juma, J. A., Mwakio, E. W., Brazeau, N., Cheruiyot, A. C., Yeda, R. A., Maraka, M. N., Okello, C. O., Kateete, D. P., Managbanag, J. R., Andagalu, B., Ogutu, B. R., & Kamau, E. (2021). *Plasmodium* interspecies interactions during a period of increasing prevalence of *Plasmodium ovale* in symptomatic individuals seeking treatment: an observational study. Lancet Microbe, 2(4), e141–e150. 10.1016/S2666-5247(21)00009-4

Akerele, D., Ljolje, D., Talundzic, E., Udhayakumar, V., & Lucchi, N. W. (2017). Molecular diagnosis of *Plasmodium ovale* by photo-induced electron transfer fluorogenic primers: PET-PCR. PLoS One, 12(6). 10.1371/journal.pone.0179178

Barber, B. E., William, T., Grigg, M. J., Yeo, T. W., & Anstey, N. M. (2013). Limitations of microscopy to differentiate *Plasmodium* species in a region co-endemic for *Plasmodium falciparum*, *Plasmodium vivax* and *Plasmodium knowlesi*. Malar. J., 12(1). 10.1186/1475-2875-12-8

Bauffe, F., Desplans, J., Fraisier, C., & Parzy, D. (2012). Real-time PCR assay for discrimination of *Plasmodium ovale curtisi* and *Plasmodium ovale wallikeri* in the Ivory Coast and in the Comoros Islands. Malar. J., 11. 10.1186/1475-2875-11-307

Bischoff, E., Guillotte, M., Mercereau-Puijalon, O., & Bonnefoy, S. (2000). A member of the *Plasmodium falciparum* Pf60 multigene family codes for a nuclear protein expressed by readthrough of an internal stop codon. Mol. Microbiol., 35(5), 1005–1016. 10.1046/j.1365-2958.2000.01788.x

Calderaro, A., Piccolo, G., Gorrini, C., Montecchini, S., Rossi, S., Medici, M. C., Chezzi, C., & Snounou, G. (2012). A New Real-Time PCR for the Detection of *Plasmodium ovale wallikeri*. PLoS One, 7(10), e48033. 10.1371/journal.pone.0048033

Calderaro, A., Piccolo, G., Perandin, F., Gorrini, C., Peruzzi, S., Zuelli, C., Ricci, L., Manca, N., Dettori, G., Chezzi, C., & Snounou, G. (2007). Genetic polymorphisms influence *Plasmodium ovale* PCR detection accuracy. J. Clin. Microbiol., 45(5), 1624–1627. 10.1128/jcm.02316-06

Chin, A. Z., Maluda, M. C. M., Jelip, J., Jeffree, M. S. bin, Culleton, R., & Ahmed, K. (2020). Malaria elimination in Malaysia and the rising threat of *Plasmodium knowlesi*. J. Physiol. Anthropol., 39(1). 10.1186/s40101-020-00247-5

Costa, G. L., Alvarenga, D. A. M. de, Assis, G. M. P. de, Aguiar, A. C. C., Louzada, J., Pereira, D. B., Pina-Costa, A. de, Hirano, Z. M. B., Moreira, S. B., Pissinatti, A., Brasil, P., Daniel-Ribeiro, C. T., Sousa, T. N. de, & Alves de Brito, C. F. (2023). Malaria mitochondrial diagnosis: challenges and pitfalls. Front. Trop. Dis., 4, 1204195. 10.3389/fitd.2023.1204195

Cox-Singh, J., Hiu, J., Lucas, S. B., Divis, P. C., Zulkarnaen, M., Chandran, P., Wong, K. T., Adem, P., Zaki, S. R., Singh, B., & Krishna, S. (2010). Severe malaria - a case of fatal *Plasmodium knowlesi* infection with post-mortem findings: a case report. Malar. J., 9(10) 10.1186/1475-2875-9-10

Deora, N., Khan, M. R., Singh, P., Zehra, N., Sircar, S., Pande, V., Mallick, P. K., & Sinha, A. (2025). Overreliance on *Plasmodium* 18S rRNA gene for malaria molecular diagnosis— inferences from systematic review. Malar. J., 24(1). 10.1186/s12936-025-05424-4

Echeverry, D. F., Deason, N. A., Davidson, J., Makuru, V., Xiao, H., Niedbalski, J., Kern, M., Russell, T. L., Burkot, T. R., Collins, F. H., & Lobo, N. F. (2016). Human malaria diagnosis using a single-step direct-PCR based on the *Plasmodium* cytochrome oxidase III gene. Malar. J., 15(1). 10.1186/s12936-016-1185-x

Fitri, L. E., Widaningrum, T., Endharti, A. T., Prabowo, M. H., Winaris, N., & Nugraha, R. Y. B. (2022). Malaria diagnostic update: From conventional to advanced method. J. Clin. Lab. Anal., 36(4), e24314. 10.1002/jcla.24314

Fornace, K. M., Topazian, H. M., Routledge, I., Asyraf, S., Jelip, J., Lindblade, K. A., Jeffree, M. S., Ruiz Cuenca, P., Bhatt, S., Ahmed, K., Ghani, A. C., & Drakeley, C. (2023). No evidence of sustained nonzoonotic *Plasmodium knowlesi* transmission in Malaysia from modelling malaria case data. Nat. Commun., 14(1), 2945. 10.1038/s41467-023-38476-8

Fuehrer, H. P., & Noedl, H. (2014). Recent advances in detection of *Plasmodium ovale*: Implications of separation into the two species *Plasmodium ovale wallikeri* and *Plasmodium ovale curtisi*. J. Clin. Microbiol., 52(2), 387–391. 10.1128/JCM.02760-13

Groger, M., Veletzky, L., Lalremruata, A., Cattaneo, C., Mischlinger, J., Zoleko, R. M., Kim, J., Klicpera, A., Meyer, E. L., Blessborn, D., Winterberg, M., Adegnika, A. A., Agnandji, S. T., Kremsner, P. G., Mordmüller, B., Mombo-Ngoma, G., Fuehrer, H. P., & Ramharter, M. (2019). Prospective Clinical and Molecular Evaluation of Potential *Plasmodium ovale curtisi* and *wallikeri* Relapses in a High-transmission Setting. Clin. Infect. Dis., 69(12), 2119–2126. 10.1093/cid/ciz131

Gujarati, S., Singal, B. R., Gupta, D., Kochar, S. K., Kochar, D. K., & Das, A. (2026). Differential expression of mitochondria-associated genes in clinical samples of *Plasmodium falciparum* showing severe manifestations. BMC Genom., 27(397). 10.1186/s12864-026-12733-8

Hayashida, K., Kajino, K., Simukoko, H., Simuunza, M., Ndebe, J., Chota, A., Namangala, B., & Sugimoto, C. (2017). Direct detection of *falciparum* and non-*falciparum* malaria DNA from a drop of blood with high sensitivity by the dried-LAMP system. Parasit. Vectors, 10(26). 10.1186/s13071-016-1949-8

He, W., Sendor, R., Potlapalli, V. R., Kashamuka, M. M., Tshefu, A. K., Phanzu, F., Kalonji, A., Ngasala, B., Thwai, K. L., Juliano, J. J., Lin, J. T., & Parr, J. B. (2024). Development of new real-time PCR assays for detection and species differentiation of *Plasmodium ovale*. PLoS Negl. Trop. Dis., 18(9): e0011759. 10.1371/journal.pntd.0011759

Hikosaka, K., Watanabe, Y. I., Kobayashi, F., Waki, S., Kita, K., & Tanabe, K. (2011). Highly conserved gene arrangement of the mitochondrial genomes of 23 *Plasmodium* species. Parasitol. Int., 60(2), 175–180. 10.1016/j.parint.2011.02.001

Isozumi, R., Fukui, M., Kaneko, A., Chan, C. W., Kawamoto, F., & Kimura, M. (2015). Improved detection of malaria cases in island settings of Vanuatu and Kenya by PCR that targets the *Plasmodium* mitochondrial cytochrome c oxidase III (cox3) gene. Parasitol. Int., 64(3), 304–308. 10.1016/j.parint.2014.09.006

Joseph, J. T., Aldritt, S. M., Unnasch, T., Puijalon, O., & Wirth, D. F. (1989). Characterization of a Conserved Extrachromosomal Element Isolated from the Avian Malarial Parasite *Plasmodium gallinaceum*. Mol. Cell. Biol., 9(9), 3621–3629. 10.1128/mcb.9.9.3621-3629.1989

Joste, V., Kamaliddin, C., Kendjo, E., Hubert, V., Argy, N., & Houzé, S. (2018). Distinction of *Plasmodium ovale wallikeri* and *Plasmodium ovale curtisi* using quantitative Polymerase Chain Reaction with High Resolution Melting revelation. Sci. Rep., 8(1). 10.1038/s41598-017-18026-1

Ke, H., Morrisey, J. M., Ganesan, S. M., Painter, H. J., Mather, M. W., & Vaidya, A. B. (2011). Variation among *Plasmodium falciparum* strains in their reliance on mitochondrial electron transport chain function. Eukaryot. Cell, 10(8), 1053–1061. 10.1128/ec.05049-11

Knowles, R., & Gupta, B. M. D. (1932). A Study of Monkey-Malaria, and Its Experimental Transmission to Man. Ind. Med. Gaz., 67(6), 301. PMID: 29010910; PMCID: PMC5231565.

Kotepui, M., Kotepui, K. U., Milanez, G. D., & Masangkay, F. R. (2020). Severity and mortality of severe *Plasmodium ovale* infection: A systematic review and meta-analysis. PLoS One, 15(6): e0235014. 10.1371/journal.pone.0235014

Lambros, C., & Vanderberg, J. P. (1979). Synchronization of *Plasmodium falciparum* erythrocytic stages in culture. J. Parasitol., 65(3), 418–420. 10.2307/3280287

Lamien-Meda, A., Fuehrer, H. P., & Noedl, H. (2019). Novel high resolution melting (HRM) and snapback assays for simultaneous detection and differentiation of *Plasmodium ovale* spp. Acta Trop., 192, 75–81. 10.1016/j.actatropica.2019.01.018

Li, J., Wirtz, R. A., Mcconkey, G. A., Sattabongkot, J., Waters, A. P., Rogers, M. J., & Mccutchan, T. F. (1995). *Plasmodium*: Genus-Conserved Primers for Species Identification and Quantitation. Exp. Parasitol., 81(2), 182–190. 10.1006/EXPR.1995.1107

Mahittikorn, A., Masangkay, F. R., Kotepui, K. U., Milanez, G. D. J., & Kotepui, M. (2021a). Comparative performance of PCR using DNA extracted from dried blood spots and whole blood samples for malaria diagnosis: a meta-analysis. Sci. Rep., 11(1), 4845. 10.1038/s41598-021-83977-5

Mahittikorn, A., Masangkay, F. R., Kotepui, K. U., Milanez, G. D. J., & Kotepui, M. (2021b). Comparison of *Plasmodium ovale curtisi* and *Plasmodium ovale wallikeri* infections by a meta-analysis approach. Sci. Rep., 11(1), 6409. 10.1038/s41598-021-85398-w

Maltha, J., Gillet, P., & Jacobs, J. (2013). Malaria rapid diagnostic tests in endemic settings. Clin. Microbiol. Infect. 19(5), 399–407. 10.1111/1469-0691.12151

Mewara, A., Sreenivasan, P., & Khurana, S. (2023). Primate malaria of human importance. Trop. Parasitol., 13(2), 73–83. 10.4103/tp.tp_79_22

Miller, R. H., Obuya, C. O., Wanja, E. W., Ogutu, B., Waitumbi, J., Luckhart, S., & Stewart, V. A. (2015). Characterization of *Plasmodium ovale curtisi* and *P. ovale wallikeri* in Western Kenya Utilizing a Novel Species-specific Real-time PCR Assay. PLoS Negl. Trop. Dis., 9(1): e0003469. 10.1371/journal.pntd.0003469

Moon, R. W., Hall, J., Rangkuti, F., Ho, Y. S., Almond, N., Mitchell, G. H., Pain, A., Holder, A. A., & Blackman, M. J. (2013). Adaptation of the genetically tractable malaria pathogen *Plasmodium knowlesi* to continuous culture in human erythrocytes. Proc. Natl. Acad. Sci. U.S.A., 110(2), 531–536. 10.1073/pnas.1216457110

Mueller, I., Zimmerman, P. A., & Reeder, J. C. (2007). *Plasmodium malariae* and *Plasmodium ovale* -the “bashful” malaria parasites. Trends Parasitol., 23(6), 278–283. 10.1016/j.pt.2007.04.009

Munyeku-Bazitama, Y., Okitale-Talunda, P., Mpingabo-Ilunga, P., Yambayamba, M. K., Tshiminyi, P. M., Umba-Phuati, A., Kimfuta, J., Phukuta, F. A., Makindu, G., Mufwaya-Nsene, R., Asari, R., Makimoto, S., Baketana, L. K., Ahuka-Mundeke, S., Isono, M., Nsio-Mbeta, J., Makiala-Mandanda, S., & Muyembe-Tamfum, J. J. (2023). High Severe Acute Respiratory Syndrome Coronavirus 2 Antibody Prevalence After the Third Epidemic Wave (May-October 2021) in Matadi, Democratic Republic of the Congo. Open Forum Infect. Dis., 10(1). 10.1093/ofid/ofad023

Oguike, M. C., Betson, M., Burke, M., Nolder, D., Stothard, J. R., Kleinschmidt, I., Proietti, C., Bousema, T., Ndounga, M., Tanabe, K., Ntege, E., Culleton, R., & Sutherland, C. J. (2011). *Plasmodium ovale curtisi* and *Plasmodium ovale wallikeri* circulate simultaneously in African communities. Int. J. Parasitol., 41(6), 677–683. 10.1016/j.ijpara.2011.01.004

Painter, H. J., Morrisey, J. M., Mather, M. W., & Vaidya, A. B. (2007). Specific role of mitochondrial electron transport in blood-stage *Plasmodium falciparum*. Nature, 446, 88–91. 10.1038/nature05572

Perandin, F., Manca, N., Calderaro, A., Piccolo, G., Galati, L., Ricci, L., Medici, M. C., Arcangeletti, M. C., Snounou, G., Dettori, G., & Chezzi, C. (2004). Development of a Real-Time PCR Assay for Detection of *Plasmodium falciparum*, *Plasmodium vivax*, and *Plasmodium ovale* for Routine Clinical Diagnosis. J. Clin. Microbiol., 42. 10.1128/jcm.42.3.1214-1219.2004

Potlapalli, V. R., Muller, M. S., Ngasala, B., Ali, I. M., Na, Y. B., Williams, D. R., Kharabora, O., Chhetri, S., Liu, M. S., Careyewend, K., Lin, F. C., Mathias, D., Tarimo, B. B., Juliano, J. J., Parr, J. B., & Lin, J. T. (2023). Real-time PCR detection of mixed *Plasmodium ovale curtisi* and *wallikeri* infections in human and mosquito hosts. PLoS Negl. Trop. Dis., 17(12), e0011274. 10.1371/journal.pntd.0011274

Preiser, P. R., Wilson, R. J. M., Moore, P. W., McCready, S., Hajibagheri, M. A. N., Blight, K. J., Strath, M., & Williamson, D. H. (1996). Recombination associated with replication of malarial mitochondrial DNA. EMBO J., 15(3), 684–693. 10.1002/j.1460-2075.1996.tb00401.x

Ramesh, S., Cihalova, D., Rajendran, E., van Dooren, G. G. & Maier, A. G. (2023). Analysis of *Plasmodium falciparum* Mitochondrial Electron Transport Chain Activity Using Seahorse XFe96 Extracellular Flux Assays. Bio-protoc. 13(21): e4863. 10.21769/BioProtoc.4863

Rojo-Marcos, G., Rubio-Muñoz, J. M., Angheben, A., Jaureguiberry, S., García-Bujalance, S., Tomasoni, L. R., Rodríguez-Valero, N., Ruiz-Giardín, J. M., Salas-Coronas, J., Cuadros-González, J., García-Rodríguez, M., Molina-Romero, I., López-Vélez, R., Gobbi, F., Calderón-Moreno, M., Martin-Echevarría, E., Elía-López, M., Llovo-Taboada, J., Lago-Nuñez, M., … Arévalo-Serrano, J. (2018). Prospective comparative multi-centre study on imported *Plasmodium ovale wallikeri* and *Plasmodium ovale curtisi* infections. Malar. J., 17(1). 10.1186/s12936-018-2544-6

Singh, B., Sung, L. K., Matusop, A., Radhakrishnan, A., Shamsul, S. S. G., Cox-Singh, J., Thomas, A., & Conway, D. J. (2004). A large focus of naturally acquired *Plasmodium knowlesi* infections in human beings. Lancet, 363(9414), 1017–1024. 10.1016/S0140-6736(04)15836-4

Snounou, G., Sharp, P. M., & Culleton, R. (2024). The two parasite species formerly known as *Plasmodium ovale*. Trends Parasitol., 40(1), 21–27. 10.1016/j.pt.2023.11.004

Snounou, S., Viriyakosol, S., Zhu, X. P., Jarra, W., Pinheiro, L., do Rosario, V. E., Thaithong, S., & Brown, K. N. (1993) High sensitivity of detection of human malaria parasites by the use of nested polymerase chain reaction. Mol. Biochem. Parasitol., 61, 315–320. 10.1016/0166-6851(93)90077-B

Stephens, J. W. W. (1922). A New Malaria Parasite of Man. Ann. Trop. Med. Parasitol., 16(4), 383–388. 10.1080/00034983.1922.11684331

Strydom, K. A., Ismail, F., & Frean, J. (2014). *Plasmodium ovale*: A case of not-so-benign tertian malaria. Malar. J., 13(85). 10.1186/1475-2875-13-85

Sutherland, C. J., Tanomsing, N., Nolder, D., Oguike, M., Jennison, C., Pukrittayakamee, S., Dolecek, C., Hien, T. T., Do Rosário, V. E., Arez, A. P., Pinto, J., Michon, P., Escalante, A. A., Nosten, F., Burke, M., Lee, R., Blaze, M., Otto, T. D., Barnwell, J. W., Pain, A., Williams, J., White, N. J., Day, N. P. J., Snounou, G., Lockhart, P. J., Chiodini, P. L., Imwong, M., & Polley, S. D. (2010). Two Nonrecombining Sympatric Forms of the Human Malaria Parasite *Plasmodium ovale* Occur Globally. J. Infect. Dis., 201(10), 1544–1550. 10.1086/652240

Syafruddin, D., Siregar, J. E., & Marzuki, S. (1999). Mutations in the cytochrome b gene of *Plasmodium berghei* conferring resistance to atovaquone. Mol. Biochem. Parasitol., 104(2), 185–194. 10.1016/S0166-6851(99)00148-6

Tamura, K., Stecher, G., & Kumar, S. (2021). MEGA11: Molecular Evolutionary Genetics Analysis Version 11. Mol. Biol. Evol., 38(7), 3022–3027. 10.1093/molbev/msab120

Tanomsing, N., Imwong, M., Sutherland, C. J., Dolecek, C., Hien, T. T., Nosten, F., Day, N. P. J., White, N. J., & Snounou, G. (2013). Genetic marker suitable for identification and genotyping of *Plasmodium ovale curtisi* and *Plasmodium ovale wallikeri*. J. Clin. Microbiol., 51(12), 4213–4216. 10.1128/jcm.01527-13

Trager, W., & Jensen, J. B. (1976). Human malaria parasites in continuous culture. Science, 193(4254), 673–675. 10.1126/science.781840

Vaidya, A. B., & Arasu, P. (1987). Tandemly arranged gene clusters of malarial parasites that are highly conserved and transcribed. Mol. Biochem. Parasitol., 22(2–3), 249–257. 10.1016/0166-6851(87)90056-9

Vaishya, S., Kumar, V., Gupta, A., Siddiqi, M. I., & Habib, S. (2016). Polypeptide release factors and stop codon recognition in the apicoplast and mitochondrion of *Plasmodium falciparum*. Mol. Microbiol., 100(6), 1080–1095. 10.1111/mmi.13369

Ye, J., Coulouris, G., Zaretskaya, I., Cutcutache, I., Rozen, S., & Madden, T. L. (2012). Primer-BLAST: A tool to design target-specific primers for polymerase chain reaction. BMC Bioinform., 13(134). 10.1186/1471-2105-13-134

Yman, V., Wandell, G., Mutemi, D. D., Miglar, A., Asghar, M., Hammar, U., Karlsson, M., Lind, I., Nordfjell, C., Rooth, I., Ngasala, B., Homann, M. V., & Färnert, A. (2019). Persistent transmission of *Plasmodium malariae* and *Plasmodium ovale* species in an area of declining *Plasmodium falciparum* transmission in Eastern Tanzania. PLoS Negl. Trop. Dis., 13(5). 10.1371/journal.pntd.0007414

